# Kir6.2-K_ATP_ channels alter glycolytic flux to modulate cortical activity, arousal, and sleep-wake homeostasis

**DOI:** 10.1101/2024.02.23.581817

**Authors:** Nicholas J. Constantino, Caitlin M. Carroll, Holden C. Williams, Carla M. Yuede, Patrick W. Sheehan, J. Andy Snipes, Erik S. Musiek, Lance A. Johnson, Shannon L. Macauley

**Affiliations:** Department of Physiology University of Kentucky, Lexington, KY; Neuroscience University of Kentucky, Lexington, KY; Sanders Brown Center on Aging, University of Kentucky, Lexington, KY; Department of Psychiatry, Wake Forest School of Medicine, Winston-Salem, North Carolina; Department of Psychiatry University School of Medicine, St Louis, MO; Department of Neurology, Washington University School of Medicine, St Louis, MO

**Author notes:** Corresponding Authors Address: Shannon L. Macauley, Associate Professor, Department of Physiology, University of Kentucky, 760 Press Avenue, Lexington, KY, 40508.

**Keywords:** K_ATP_ channels, metabolism, excitability, sleep, arousal, EEG, behavior, metabolomics

## Abstract

Metabolism plays an important role in the maintenance of vigilance states (e.g. wake, NREM, and REM). Brain lactate fluctuations are a biomarker of sleep. Increased interstitial fluid (ISF) lactate levels are necessary for arousal and wake-associated behaviors, while decreased ISF lactate is required for sleep. ATP-sensitive potassium (K_ATP_) channels couple glucose-lactate metabolism with neuronal excitability. Therefore, we explored how deletion of neuronal K_ATP_ channel activity (Kir6.2-/- mice) affected the relationship between glycolytic flux, neuronal activity, and sleep/wake homeostasis. Kir6.2-/- mice shunt glucose towards glycolysis, reduce neurotransmitter synthesis, dampen cortical EEG activity, and decrease arousal. Kir6.2-/- mice spent more time awake at the onset of the light period due to altered ISF lactate dynamics. Together, we show that Kir6.2-K_ATP_ channels act as metabolic sensors to gate arousal by maintaining the metabolic stability of each vigilance state and providing the metabolic flexibility to transition between states.

**Highlights:**

- Glycolytic flux is necessary for neurotransmitter synthesis. In its absence, neuronal activity is compromised causing changes in arousal and vigilance states despite sufficient energy availability.
- With Kir6.2-K_ATP_ channel deficiency, the ability to both maintain and shift between different vigilance states is compromised due to changes in glucose utilization.
- Kir6.2-K_ATP_ channels are metabolic sensors under circadian control that gate arousal and sleep/wake transitions.

**Graphical Abstract:** 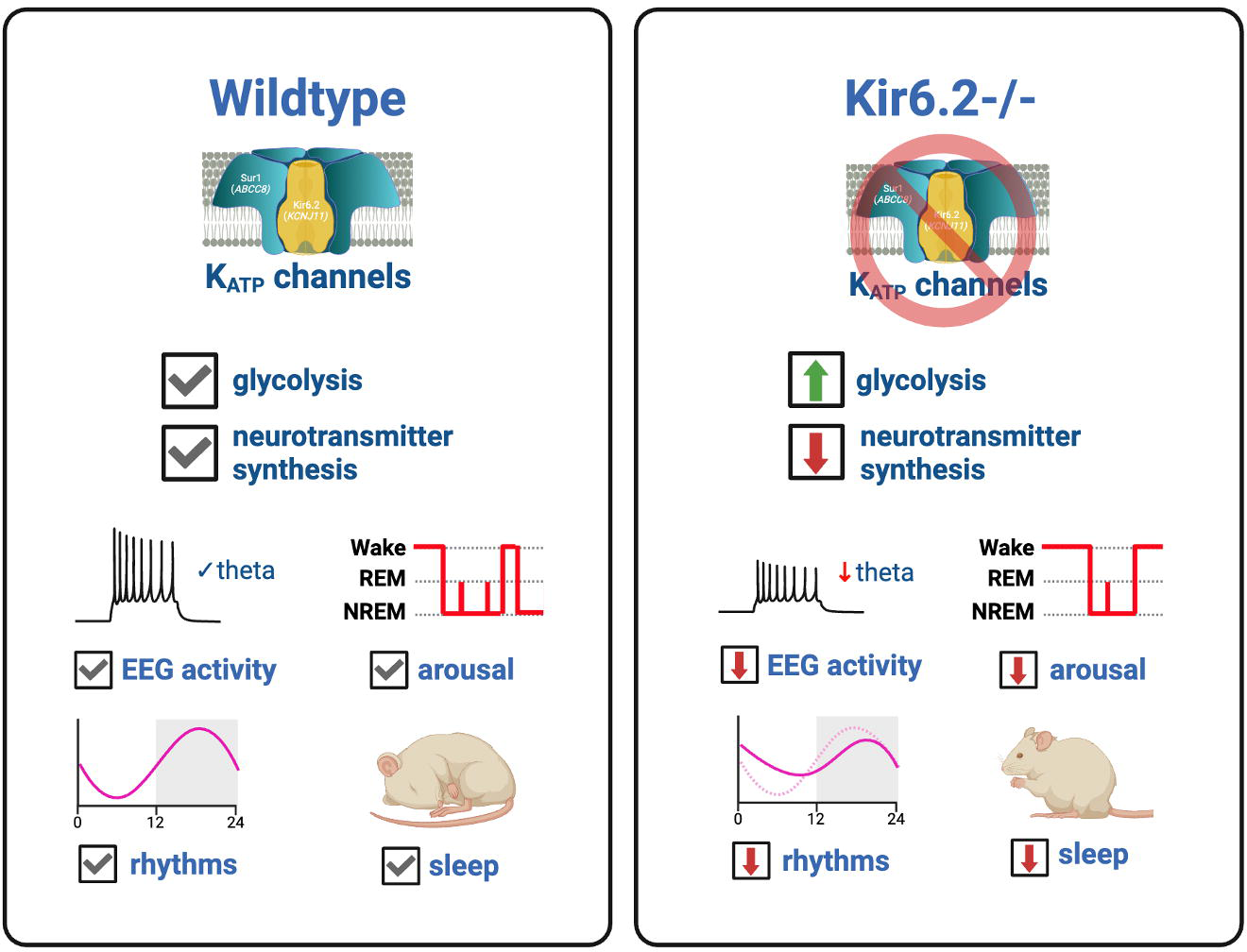

## Introduction

Metabolism is intrinsically and bidirectionally linked to vigilance states (e.g. wake, NREM, REM) and arousal (1–3). Traditionally, it was thought that heightened neuronal activity and wake-associated behaviors have the greatest impact on cerebral metabolism, but recent evidence suggests that sleep requires its own distinct metabolic signature(1, 2). Brain lactate levels are a biomarker of sleep/wake states. Brain interstitial fluid (ISF) lactate rapidly rises with the onset of wake and quickly drops with the onset of NREM sleep(3–6). The metabolic rate of glucose changes across vigilance states, where heightened arousal and complex behavioral states are more strongly associated with lactate production than oxidative metabolism (1, 2, 7). Conversely, ISF lactate levels must drop to sustain NREM sleep, quiet wakefulness, or decreased arousal(6). Cerebral metabolism is not only affected by vigilance states, but also by circadian rhythms(8). Brain glucose metabolism increases during the active period and decreases during the inactive period when animals typically sleep. While sleep/wake cycles and circadian rhythms are interconnected, ISF lactate fluctuations are more closely associated with sleep/wake transitions than ISF glucose (6). Taken together, this demonstrates that changes in cellular metabolism can regulate vigilance states and arousal; however, many of the molecular mechanisms governing the relationship between metabolism and vigilance states are not well defined.

One mechanism that couples metabolism with neuronal activity are ATP-sensitive potassium (K_ATP_) channels. When ATP levels are high, K_ATP_ channels close, biasing cells towards excitability.

Conversely, when ATP levels are low, the channels remain open, hyperpolarizing the cell, and limiting excitability. K_ATP_ channels are composed of 4 pore forming subunits (e.g. Kir6.1, Kir6.2) and 4 sulfonylurea regulatory receptors (e.g. Sur1, Sur2A/B). K_ATP_ channels containing the Kir6.2 subunit, or Kir6.2-K_ATP_ channels, are found on excitatory and inhibitory neurons within the brain, where they act as metabolic sensors and contribute to excitatory/inhibitory balance(9, 10). Treatment with K_ATP_ channel antagonists (e.g. sulfonylureas) increase neuronal firing in Kir6.2-K_ATP_ channel expressing neurons, while K_ATP_ channel agonists (e.g. diazoxide) reduce excitability. Furthermore, during peripheral hyperglycemia, Kir6.2-K_ATP_ channels couple elevations in ISF glucose levels with ISF lactate levels to sustain neuronal firing (6, 10). However, ISF glucose and lactate metabolism are uncoupled in mice lacking Kir6.2-K_ATP_ channels (e.g. Kir6.2-/- mice), yet the impact on excitability and behavior is largely unknown(10). Because K_ATP_ channels are found on both glucose sensing(11) and orexinergic neurons (12) in the hypothalamus(13), they are likely involved in sleep/wake homeostasis, circadian rhythms, and arousal(7, 14–18).

While the loss of Kir6.2-K_ATP_ channels (e.g. Kir6.2-/- mouse) result in altered ISF glucose-lactate coupling(10), it is unknown how deleting this metabolic sensor affects the brain’s glycolytic flux, neuronal activity, vigilance states, or behavior. Since Kir6.2-K_ATP_ channels are necessary for glucose- dependent increases in ISF lactate, we hypothesized that the deletion of these channels would dampen neuronal activity and disrupt the brain’s diurnal and sleep-wake rhythms. Therefore, we explored how Kir6.2-K_ATP_ channel deletion using Kir6.2-/- mice impacted the relationship between metabolism, excitability, biological rhythms, and sleep (Figure 1A). We used stable isotope resolved metabolomics (SIRM) to explore U-^13^C-glucose utilization in the brain as a consequence of K_ATP_ channel deletion.

**Figure 1.**
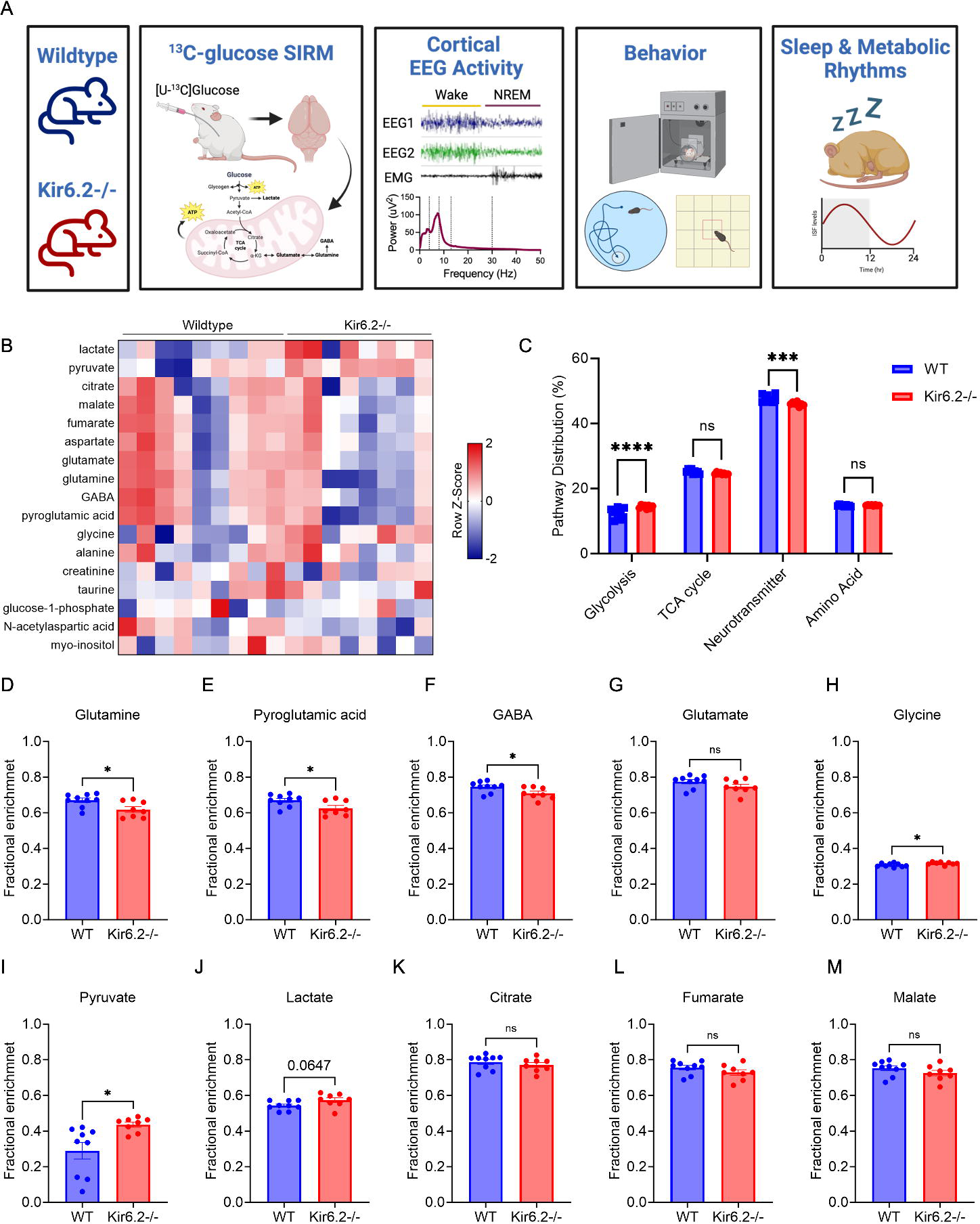
Kir6.2 -/- brains shunt glucose towards glycolysis and away from neurotransmitter synthesis. (A) Schematic of experimental design. (B) Heatmap representation of detectable metabolites following ^13^C-glucose administration. (C) Incorporation of ^13^C-glucose into metabolic pathways. Kir6.2 -/- mice shunt more glucose towards glycolysis and away from neurotransmitter synthesis (p<0.0001 and p=0.0003, respectively). (D-F) Glutamine, pyroglutamic acid, and GABA fractional enrichment is decreased in Kir6.2-/- mice (p<0.05). (G) Glutamate fractional enrichment is unaltered in Kir6.2-/- mice. (H-I) Glycine and pyruvate fractional enrichment is increased in Kir6.2-/- mice p<0.05). (J) Lactate fractional enrichment is trends towards an increase in Kir6.2-/- mice (p=0.0647). (K-M) Citrate, fumarate, and malate fractional enrichment is unaltered in Kir6.2-/- mice. Data reported as ± SEM. n = 8-9 mice/genotype. Pathway distribution significance determined by 2-way ANOVA with Turkey multiple comparisons for post hoc analysis. Fractional enrichment significance determined via unpaired t-test. *p<0.05, ***p<0.001, ****p<0.0001.

Next, we used cortical EEG/EMG to investigate how Kir6.2-K_ATP_ channel deletion impacts EEG power and its relationship with arousal, anxiety, and cognition. Finally, using simultaneous EEG/EMG recordings with intracranial monitoring of ISF glucose and lactate levels, we explored how K_ATP_ channel deletion affected sleep/wake homeostasis and metabolic rhythms. Together, our data shows Kir6.2-K_ATP_ channels act as metabolic sensors to gate glycolytic flux and regulate cortical EEG activity, arousal, and sleep/wake homeostasis. Kir6.2-K_ATP_ channel activity allows neurons to sense changes in energy availability, which then impacts their ability to both maintain and shift between different vigilance states and regulate neuronal excitability. More broadly, these studies highlight the importance of metabolic flexibility to maintain arousal, biological rhythms, and sleep/wake homeostasis.

## Results

### Kir6.2-K_ATP_ deficient (Kir6.2-/-) brains increase glycolysis at the expense of neurotransmitter synthesis

Following oral gavage of U-^13^C-glucose, cerebral glucose metabolism was quantified via stable isotope resolved metabolomics (SIRM)(19). A heatmap representation of relative abundance for each metabolite in Kir6.2-/- and WT brains is shown in Figure 1B. ^13^C-labeled metabolites were binned into pathways to explore patterns of glucose utilization more broadly in Kir6.2-/- mice. Differences in neurotransmitter synthesis (glutamate, glutamine, GABA, pyroglutamic acid, NAA, myo-inositol, aspartate) and glycolysis (pyruvate, lactate, glucose-1-phosphate) were observed in Kir6.2-/- mice compared to WT (Fig 1C, p<0.0005, p<0.0001), while metabolites associated with the TCA cycle (citrate, malate, fumarate) and amino acid synthesis (alanine, creatinine, taurine, glycine) were similar across groups. Kir6.2-/- mice had increased pyruvate and lactate synthesis (Fig 1I-J, p<0.05, p=0.0647), while the abundance of TCA cycle intermediates, citrate, fumarate, and malate, were unaltered in Kir6.2-/- mice compared to WT (Fig 1K-M). Kir6.2-/- mice had decreased ^13^C-labeled glutamine and pyroglutamic acid (Fig 1D-E, p<0.05), precursors to neurotransmitters glutamate and GABA. GABA synthesis was decreased in Kir6.2-/- mice (Fig 1F, p<0.05), while glutamate remained unchanged (Fig 1G). Glycine, an amino acid that can also acts an inhibitory neurotransmitter and co-agonist with glutamate for NMDA receptor-mediated excitatory neurotransmission, increased in Kir6.2-/- mice compared to WT mice (Fig 1H, p<0.05). Interestingly, the total abundance of all brain metabolites was similar across groups (Fig S1), suggesting that shifts in glucose utilization are not due to decreased energy availability. This suggests that neuronal excitability and the excitatory-inhibitory balance are potentially compromised in Kir6.2-/- mice due to increased glycolysis.

### Kir6.2-K_ATP_ channel deletion dampens absolute EEG power, reduces alpha and theta activity, and causes behavioral impairments associated with arousal, anxiety, and cognition

Based on results from SIRM data suggesting Kir6.2-/- mice have alterations in neurotransmitter synthesis, we explored whether Kir6.2-/- mice have changes in neuronal activity and behavior. Cortical activity was recorded using skull screw EEGs in Kir6.2-/- and WT mice. We found that absolute power, a measure of the summative strength of cortical EEG activity, was dampened in Kir6.2-/- mice across all frequencies and sleep-wake states (Fig 2A-C, p<0.0001). This demonstrates that changes in metabolic flux in Kir6.2-/- mice can also decrease EEG power.

**Figure 2.**
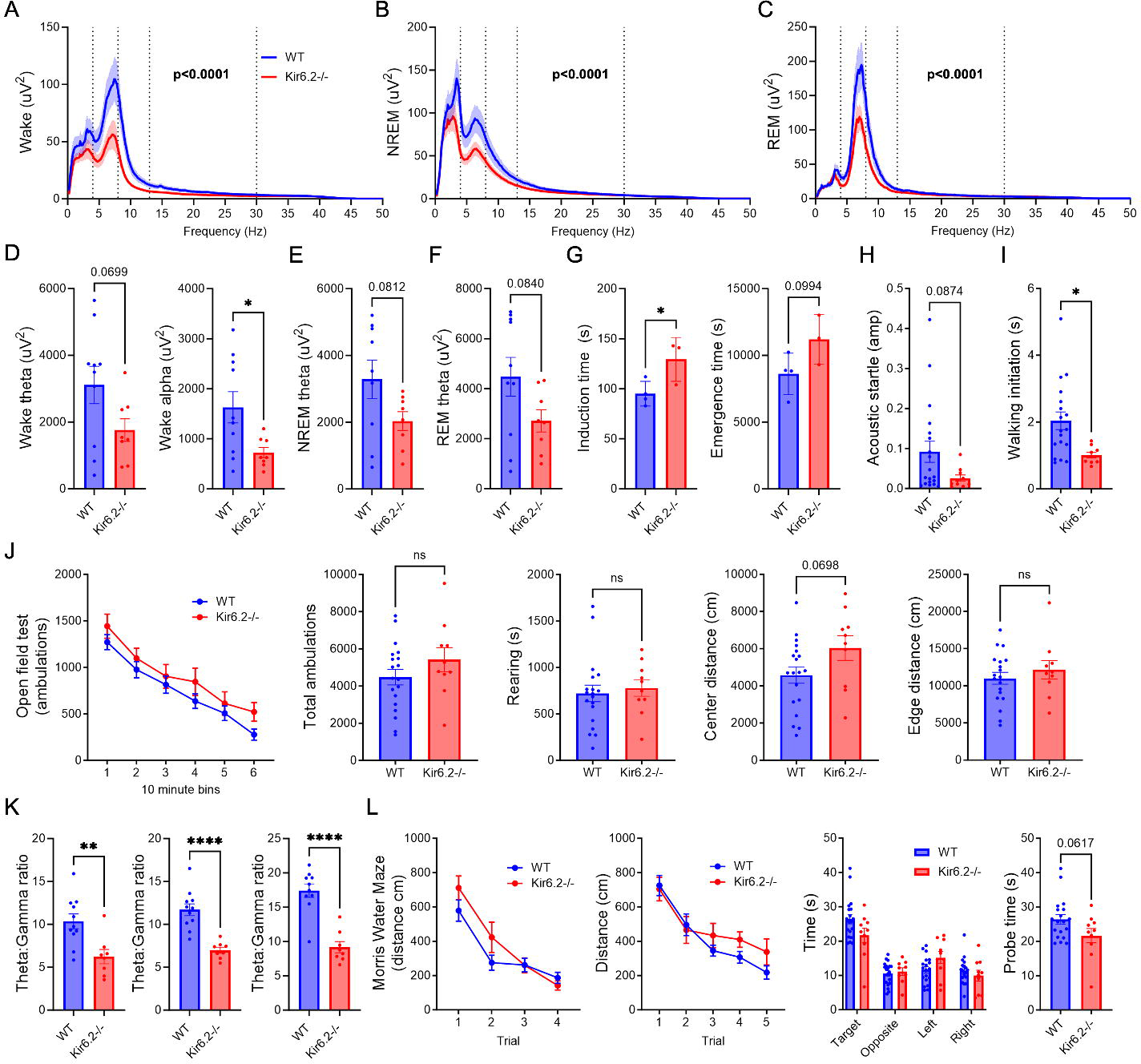
Reductions in absolute cortical EEG power across sleep-wake states is reflected in changes in arousal, anxiety, and cognitive behaviors. (A-C) Total cortical EEG absolute power is decreased across wake, NREM and REM in Kir6.2-/- mice (genotype x frequency, p<0.0001). (D) In wake, absolute theta power trends towards a decrease (p=0.0699) and alpha power is decreased (p<0.05) in Kir6.2-/- mice. (E-F) In NREM and REM, absolute theta power trends towards a decrease (p<0.0840) in Kir6.2-/- mice. (G) Kir6.2-/- mice trend towards a reduced behavioral response to acoustic startle (p=0.0874). (H) In a walking initiation task, Kir6.2-/- mice showed a decreased latency to explore a novel environment (p<0.05). (I) Kir6.2-/- mice have similar levels of general activity and ambulations compared to WT mice. However, Kir6.2-/- mice trend towards more distance traveled in the center of the activity chamber (p=0.0698). (J) Relative theta:gamma ratio, an EEG marker of cognitive function, is decreased in Kir6.2-/- mice across wake, NREM and REM sleep (p<0.01, p<0.0001). (K) Distance traveled to find visible and submerged platform in Morris water maze test (MWM) decreases with trials at similar rates in Kir6.2-/- and WT mice. There is a quadrant x genotype interaction for spatial bias (p=0.0168). Kir6.2-/- mice trend towards spending less time in the target quadrant, indicated by probe time, suggesting memory impairment (p=0.0617). Data reported as means ± SEM. n = 8-11 mice/genotype for EEG/EMG and n = 10-20 mice/genotype for behavior. Total absolute power significance determine by 2-way ANOVA with Sidak multiple comparisons correction for post hoc analysis. Individual frequency band and theta:gamma ratio significance determined by unpaired t-test. Significance of distance traveled, spatial bias and ambulation’s determined via a 2-way ANOVA with Bonferroni multiple comparison’s correction for post hoc analysis. All other significance determined via unpaired t-tests. *p<0.05, **p<0.01, ****p<0.0001

Absolute power was then binned into specific frequency bands, including delta (0.5-4Hz), theta (4-8 Hz), alpha (8-13 Hz), beta (13-30Hz), and gamma (30-50Hz), to explore whether changes in absolute EEG power could predict alterations in behavior. We found that theta (4-8 Hz) and alpha (8-13 Hz) power- frequency bands typically associated with arousal, sleep pressure, and state switching (20–24)- were decreased in Kir6.2 -/- mice during wake (Fig 2D, p=0.0699, p<0.05). Theta band activity, which builds during NREM sleep and dominates EEG activity during REM sleep(24–27), also trended towards decreased power in both NREM and REM in Kir6.2-/- mice (Fig 2E-F, p=0.0812, p=0.0840).

Given that alpha and theta bands were reduced with Kir6.2-K_ATP_ channel deletion, we explored whether behaviors associated with arousal were affected in Kir6.2-/- mice. First, we explored whether Kir6.2-/- mice responded differently to anesthesia. We used this test as a proxy for how rapidly mice transition between consciousness or vigilance states (Figure 2G). Induction time (e.g. wake to anesthetization) and emergence time (e.g. anesthetization to wake) were both longer for Kir6.2-/- mice compared to WT mice (p<0.05 and p<0.09, respectively), suggesting transitions between vigilance states were delayed. On the acoustic startle test, Kir6.2-/- mice trended towards a decreased startle response compared to controls (Fig 2G, p=0.0874), further suggesting alterations in arousal. Changes in theta activity are not only linked with arousal, but also anxiety(28, 29). Therefore, we explored anxiety-like behaviors in Kir6.2-/- mice through a walking initiation task and locomotor activity. Kir6.2-/- mice are quicker to investigate a novel context compared to WT on a walking initiation task (Fig 5H, p<0.05), suggesting decreased anxiety-like behaviors. While no overt motor impairments were observed in Kir6.2-/- mice during an open field test, Kir6.2-/- mice explored the center of the cage more than WT mice (Fig 5I, p=0.0698), which is also indicative of an anxiolytic phenotype. Taken together, decreased alpha and theta activity are associated with decreased arousal and anxiety-like behaviors in the Kir6.2-/- mice.

Since Kir6.2-K_ATP_ channel deletion impacts theta activity, we also quantified the relative ratio of theta to gamma power, a measure of memory and cognitive control, in Kir6.2-/- and WT mice. In the Kir6.2-/- mice, theta:gamma ratio was decreased across wake, NREM, and REM (Fig 2J, p<0.01, p<0.0001), suggesting Kir6.2-/- mice may suffer from memory impairment(30, 31). Therefore, we performed the Morris water maze (MWM), a test of cognitive function. While Kir6.2-/- mice were able to learn this task at similar rates to WT mice, time spent in the target quadrant during the probe trial was reduced (Fig 2K, p<0.0617), which is indicative of memory and cognitive impairments(32).

### Kir6.2-K_ATP_ channel deletion delays wake-to-sleep transitions, reduces sleep time, and shifts relative power towards high frequency activity

Since alpha and theta activity are strongly associated with arousal and sleep pressure, we explored whether Kir6.2-/- mice have impairments in sleep/wake cycles across the circadian day. Sleep staging of EEG/EMG records was performed and average minutes per hour of wake, NREM, and REM were calculated for Kir6.2-/- and WT mice. Over the 24-hour day, there were modest alterations in sleep and wake time in Kir6.2-/- mice, specifically at ZT0-3, or dark-to-light transitions, when mice should spend more time asleep (Fig 3A-C). At ZT3, time spent in wake was increased (p<0.05), while both NREM (p<0.05) and REM (p<0.01) sleep were decreased. This suggests a delayed transition to sleep in Kir6.2-/- mice.

**Figure 3.**
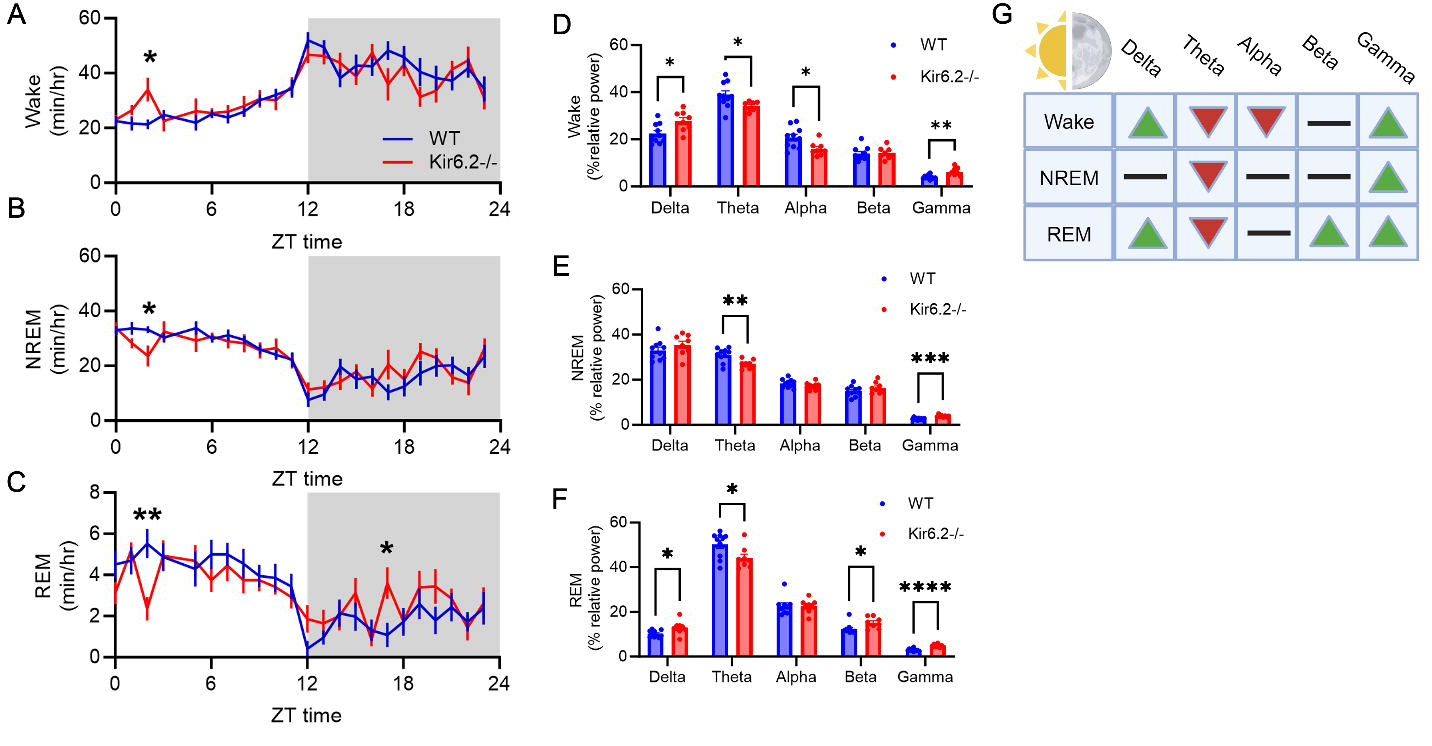
Kir6.2-/- mice have reduced sleep time and a relative shift in EEG power from theta to gamma across all vigilance states. (A-C) Minutes per hour spent in wake, NREM, and REM across the diurnal day. Kir6.2-/- mice have increased wake (p<0.05), decreased NREM sleep (p<0.05), and decreased REM at ZT3 (p<0.01). Kir6.2-/- mice also have increased REM at ZT17(p<0.05). (D) Kir6.2-/- mice lose theta and alpha power (p<0.05) and gain delta and gamma power (p<0.05, p<0.01) during wake. (E) Kir6.2 -/- mice lose theta and gain gamma power during NREM (p<0.01, p<0.001). (F) Kir6.2- /- lose theta power (p<0.05) and gain delta, beta, and gamma power (p<0.05, p<0.05, p<0.0001) during REM. (G) Summary table. Data reported as means ± SEM. n = 8-11 mice/genotype for EEG analysis and n = 2 WT mice for JTK analysis. 24 hour minute/hour sleep-wake state and relative power significance determined by 2-way ANOVA with Turkey multiple comparisons correction for post hoc analysis. *p<0.05,**p<0.01,***p<0.001, ****p<0.0001.

Next, we quantified the distribution of relative EEG power across sleep/wake states to assess the quality of wake, NREM, and REM vigilance states (Fig 3D-G). During wake, there was increased relative delta and gamma power (Fig 3D, p<0.05, p<0.01), with a corresponding decrease in theta and alpha power in Kir6.2-/- mice (Fig 3D, p<0.05). Shifts towards increased delta activity during wake suggest either increased homeostatic sleep drive or quiet wakefulness(33–36), which are rarely accompanied by increased gamma activity since gamma is associated with heightened arousal and complex cognitive processing. In NREM sleep, Kir6.2-/- mice had decreased relative theta power (Fig 3E, p<0.01), with a concurrent shift towards increased gamma (Fig 3E, p<0.001). Theta power should build over NREM bouts to drive REM sleep, suggesting an inability to switch between vigilance states. NREM sleep is traditionally composed of low frequency delta activity so increased high frequency gamma activity, known to drive depolarization or UP states(37), would decrease the quality of NREM sleep. In REM, Kir6.2-/- had increased relative delta, beta, and gamma power (Fig 3F, p<0.05, p<0.0001), with a decrease in relative theta power (p<0.05). This suggests that the integrity of REM sleep is also compromised in Kir6.2-/- mice since theta power should dominate REM sleep(38). Across all vigilance states, a shift from decreased theta and increased gamma power is observed, suggesting an uncoupling of theta:gamma EEG activity and tightly coordinated neuronal activity.

Since most sleep/wake changes occurred during the early stages of the light period, we explored if sleep/wake and relative power changes were specific to the light period (ZT0-12). We found that there is a trend for increased time spent in wake in Kir6.2-/- mice (Fig S2A, p=0.0709) during the light period, but not in the dark period (Fig S2B-F). This suggests that changes in sleep/wake homeostasis during the light period drive changes seen in Kir6.2-/- mice across the 24-hour day.

Given that changes in sleep/wake cycles occurred at specific times of day, we explored whether gene expression of Kir6.2-K_ATP_ channels was rhythmic. Kir6.2-K_ATP_ channels are heteroctameric and composed of Kir6.2 subunits and Sur1 sulfonylurea binding sites. The genes, *Kcnj11* and *Abcc8*, code for the Kir6.2 and Sur1 proteins, respectively. To assess gene rhythmicity, wildtype mice were euthanized every 2 hours over a 24-hour time period and Jonckheere–Terpstra–Kendall (JTK) analysis of *Kcnj11* and *Abcc8* gene expression was quantified (data not shown). Both *Kcnj11* and *Abcc8* are rhythmic, with peak expression during the light period at ZT10-12, then decreasing during the dark period or active phase. This suggests Kir6.2-K_ATP_ channel expression is under circadian control, which may provide one explanation for why the changes in excitability and sleep/wake states are more pronounced at specific times of day.

### Kir6.2-K_ATP_ channel deletion causes phase shifts in diurnal rhythms of ISF lactate, which corresponds with delays in sleep-wake transitions

Following our EEG data suggesting alterations in state switching in Kir6.2-/- mice, we analyzed whether there were alterations in ISF lactate, a metabolic biomarker of sleep/wake transitions (Figure 4A)(3, 4, 39), during NREM-to-wake and wake-to-NREM transitions. We found that as Kir6.2-/- mice transition from sleep to wake, ISF lactate does not increase as it does in WT mice (Fig 4B, p<0.05). When examining wake-to-NREM transitions, Kir6.2 -/- mice did not display subsequent decreases in ISF lactate compared to WT as they go to sleep (Fig 4B, p<0.05). When investigating ISF lactate and ISF glucose fluctuations across the day, ISF lactate levels positively correlated with wake in both Kir6.2-/- and WT mice (Fig S3A, p<0.0001). Again, suggesting that metabolism does fluctuate with sleep/wake states. In NREM and REM, the opposite pattern was observed. In both Kir6.2-/- and WT mice, ISF lactate levels were negatively correlated with time spent in sleep (Fig S3B-C, p<0.0001). Interestingly, Kir6.2-/- mice lose all correlations between sleep-wake states and ISF glucose that are present in WT mice (Fig S3D-F, p=0.0006). Together, these data suggest that Kir6.2-/- mice lack the lactate trigger, or metabolic flexibility, to switch between vigilance states because of the alterations in glucose utilization.

**Figure 4.**
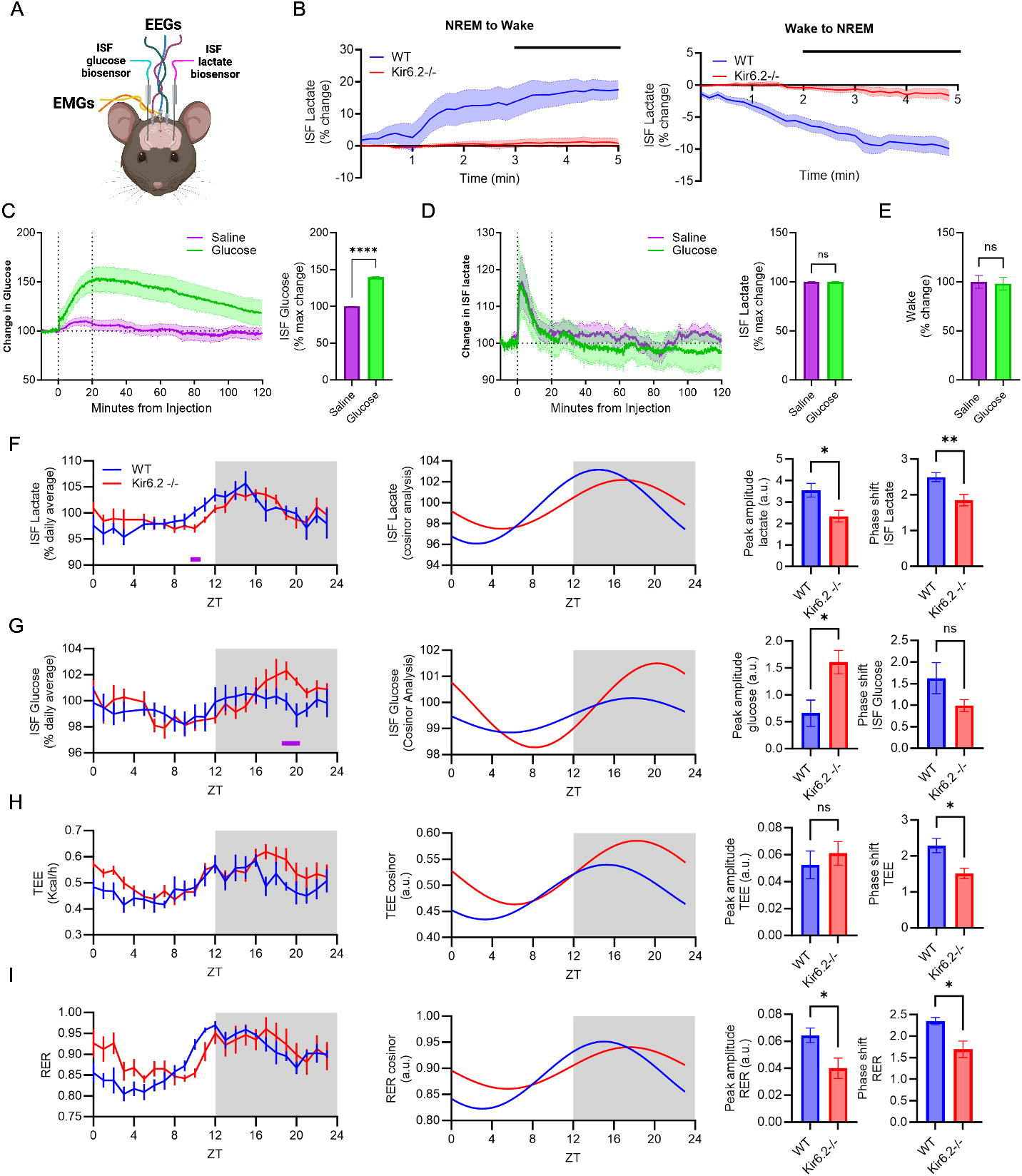
Brain and peripheral metabolic rhythms change with Kir6.2-K_ATP_ channel deletion. (A) Schematic of EEG, EMG, and biosensor setup in mice. (B) ISF lactate does not increase during NREM to wake transitions in Kir6.2-/- mice (p<0.05). Nor, does ISF lactate decrease during wake to NREM sleep transitions in Kir6.2-/- mice (p<0.05). (C) Ip glucose challenge increases ISF glucose in Kir6.2-/- mice (p<0.0001). (D) IP glucose challenge does not increase in ISF lactate in Kir6.2-/- mice. (E) % change in time spent in wake following an ip glucose injection is unchanged compared to saline injections in Kir6.2-/- mice. (F) The rise in ISF lactate preceding the light-dark switch is delayed in the Kir6.2-/- mice (p<0.05). Cosinor analysis of ISF lactate rhythms show decreased amplitude (p<0.05) and a phase shift in diurnal rhythms in Kir6.2-/- mice (p<0.001). (G) During the dark phase, the peak in ISF glucose levels is increased in Kir6.2-/- mice at ZT19-20 (p<0.05). Cosinor analysis of ISF glucose shows increased amplitude (p<0.05) and a greater peak in the dark period in in Kir6.2-/- mice. (H) Fluctuations in total energy expenditure (TEE) is phase shifted in Kir6.2-/- mice compared to WT mice (time X genotype interaction, p= 0.0005). (I) A phase shift and decreased amplitude in respiratory exchange ratio (RER) is present in Kir6.2-/- mice (time X genotype interaction, p<0.0001). Data reported as means ± SEM. Sleep wake transitions % change in ISF lactate (n= 8-18 mice/genotype). Significance determined by 2-way ANOVA with Sidak multiple comparisons correction for post hoc analysis. % change from baseline injection data determined by one-way ANOVA with Dunnett multiple comparisons correction for post hoc analysis (n=8 Kir6.2-/- mice). Diurnal rhythms of ISF lactate, ISF glucose, TEE and RER determined via 2-way ANOVA with Sidak or Turkey multiple comparisons post hoc analysis. ISF glucose and lactate n=8-11 mice/genotype. TEE and RER n=5 mice/genotype. *p<0.05, **p<0.01.

Since a peripheral hyperglycemic challenge is sufficient to stimulate wake through ISF lactate production(6), we examined whether Kir6.2-/- mice reacted to a hyperglycemic challenge by increasing ISF lactate levels and time spent awake. Following a hyperglycemic challenge, we observed increased ISF glucose (Fig 4C, p<0.0001). However, neither ISF lactate levels nor time spent awake in Kir6.2-/- mice increased as it should in response to a hyperglycemic challenge (Fig 4D-E). This shows that Kir6.2-K_ATP_ channels are necessary to couple changes in metabolism with changes in arousal.

Since K_ATP_ channels couple metabolism and excitability and gene expression of Kir6.2-K_ATP_ channels is rhythmic, we next asked how changes in the interstitial fluid (ISF) levels of glucose and lactate changed over the 24-hour day. ISF lactate fluctuations are rhythmic (Supplemental Table 1), peaking during the dark period in both WT and Kir6.2-/- mice (Fig 4F). Kir6.2-/- mice have lower levels of ISF lactate at ZT10 with a delay to rise at the light-dark transition (Fig 4F, p<0.05). Both JTK and cosinor analyses show that Kir6.2-/- mice have a lower peak amplitude of ISF lactate and a phase shift in diurnal rhythms compared to controls (Fig 4F, p<0.05, p<0.01 & Supplemental Table 1). ISF glucose levels peak in the dark period for both groups, with Kir6.2-/- mice having a greater peak at ZT19-20 (Fig 4G, p<0.05 & Supplemental Table 1).

Next, we explored whether alterations in metabolic rhythms were restricted to the brain in Kir6.2-/- mice. We found that the diurnal shifts in cerebral metabolic rhythms were mirrored in the periphery and correlated with peripheral metabolic fluctuations (Fig 4H-I, Fig S4-5, Supplemental Table 1). In order to characterize peripheral metabolism over the diurnal day, Kir6.2-/- and WT mice underwent indirect calorimetry(40). Both total energy expenditure (TEE) and respiratory exchange ratio (RER) were rhythmic in Kir6.2-/- and WT mice (Supplemental Table 1). However, Kir6.2-/- mice had shifted in TEE and RER rhythms compared to WT mice, with a time x genotype interaction (Fig 4H-I, p<0.0005). Both TEE and RER were strongly correlated with time spent in wake, NREM, and REM (Fig S4). Interestingly, TEE rhythms reflected increases in wake during the first few hours of the light period and elevated brain glucose levels during the dark period. TEE was correlated with ISF lactate in both

Kir6.2-/- and WT mice (Fig S5A, p<0.0001), but the relationship between TEE and ISF glucose was lost in Kir6.2-/- mice (Fig S5C). Additionally, RER rhythms largely correlated with ISF lactate and glucose rhythms (Fig S5 B&D), where an increase in RER during the first few hours of the light period indicated increased utilization of carbohydrates as a fuel source and a delay in fuel switching towards fatty acids during sleep. Furthermore, RER shows a similar phase shift to ISF lactate rhythms in Kir6.2-/- mice, with a delay to rise around the light-to-dark transition, illustrating the animal’s shift in arousal (Fig 4I, p<0.05). Overall, this illustrates that changes in central and peripheral metabolism are tightly coupled, with TEE and RER correlating strongly with ISF lactate fluctuations.

## Discussion

Healthy brain function relies on the dynamic interplay between metabolism and excitability, where different vigilance states have their own metabolic and activity-dependent profiles. The ability of the brain to execute complex behavior or conversely, engage in restorative states, relies on coordinated changes in metabolism and neuronal excitability. While it is well established that patterns of neuronal activity underly changes in behavior, less is known about how metabolic flexibility and glycolytic flux can respond to and drive changes in arousal. When considering glucose metabolism as a contributor to vigilance states, our work demonstrates that glucose must be considered as both a fuel source for ATP generation, but also as a substrate for nonoxidative metabolism, biosynthesis, and redox regulation.

Our data suggests alterations in glycolytic flux have profound effects on cellular function, demonstrating its importance for the stability and flexibility of vigilance states.

^13^C-glucose SIRM experiments illustrated that Kir6.2-K_ATP_ channel deletion alters glycolytic flux where glucose is shunted towards glycolysis at the expense of neurotransmitter synthesis. It is notable that the change in glycolytic flux did not occur as a result of an overall ATP shortage, but as a reaction to the way neurons detect and use glucose as a substrate for biosynthesis and specific physiological processes. This subtle alteration in glycolytic flux was sufficient to decrease cortical EEG power, the integrity of sleep/wake states, and the flexibility to transition between wake, NREM, and REM. We found that absolute power was dampened across all sleep/wake states in Kir6.2-/- mice, with a shift in relative power from lower (e.g. theta and alpha) to higher (e.g. gamma) frequency bands. The loss of theta power is interesting since this was observed in wake, NREM, and REM. The likelihood of transitioning between vigilance states, be it wake-to-NREM or NREM-to-REM, appears to be driven by increased theta power(41, 42). During wake, the ability to attend, perform, and sustain different levels of arousal depends upon theta activity in the anterior cortex (43). Therefore, this study identifies a new role for Kir6.2-K_ATP_ channels in regulating behaviors associated with theta power. Furthermore, alpha power is associated with drowsiness or quiet wakefulness, which is necessary for the gradual transition to sleep. The reduction of alpha power during wake reinforces the idea that Kir6.2-K_ATP_ channels are important for modulating arousal and the state-shift between sleep and wake. Our conclusions from animal studies are further supported by clinical studies where individuals with *KCNJ11* mutations report higher incidences of problems with sleep, anxiety, and attention(18).

The behavioral manifestations of Kir6.2-K_ATP_ channel deficiency were nuanced. Ultimately, whether looking at arousal, anxiety, cognition, or sleep, mice lacking K_ATP_ channel activity largely retained the ability to complete tasks or maintain sleep/wake homeostasis. However, the robustness of a specific behavior or the ease to switch between different vigilance states was compromised in Kir6.2-/- mice. This suggested that Kir6.2-K_ATP_ channels are important regulators of metabolic flexibility and cognitive processes like arousal. Again, this highlights the idea that the metabolic flexibility is as important as total energy availability.

Another key observation from this study is that Kir6.2-K_ATP_ channels are necessary for glucose-lactate coupling, which is integral for sleep/wake transitions. Brain, or ISF, lactate levels are a biomarker for sleep. Increased lactate triggers wake, while decreased ISF lactate levels are necessary to transition to NREM sleep. Our ^13^C-glucose experiments show that: 1) the relative production of lactate in the Kir6.2-/- brain is higher than WT, 2) the total abundance of brain lactate is comparable between Kir6.2-/- mice and WT, and 3) yet the pool of ISF lactate is reduced or less dynamic in Kir6.2-/- mice compared to WT. This demonstrates that Kir6.2-K_ATP_ channels not only act as metabolic sensors to couple metabolism with excitability, but also regulate the extracellular flux of lactate which has pleiotropic effects on cortical activity and behavior. Herein, we demonstrate that alterations in lactate dynamics due to Kir6.2-K_ATP_ channel deficiency underly the integrity of and transition between vigilance states. Phase delays in ISF lactate accompany disruptions in sleep homeostasis, most notably in the ability for mice to transition from wake-to-sleep. While it is known that K_ATP_ channels are localized to the hypothalamus and play a role in orexinergic neuronal functioning, we are the first to show that *Kcnj11* and *Abcc8*, the genes regulating Kir6.2-/- and Sur1 expression, are rhythmic. This shows an interesting interplay between sleep/wake cycles and circadian rhythms, where K_ATP_ channels may serve as a metabolic zeitgeber for circadian rhythms and sleep/wake cycles.

Taken together, our studies describe how Kir6.2-K_ATP_ channels play an important role in glycolytic flux, where glucose serves as both a metabolite and biosynthetic substrate for the brain. Kir6.2-K_ATP_ channel deficiency causes subtle changes in glycolytic flux that alter cortical EEG activity, leading to deficits in arousal, attention, and sleep/wake architecture. This highlights the important relationship between metabolic flexibility and the maintenance of different vigilance states. Ultimately, this has repercussions for many diseases including, type-2-diabetes, developmental delay, epilepsy and neonatal diabetes (DEND syndrome), and Alzheimer’s disease(10, 18), where K_ATP_ channels are known to play a role in disease pathogenesis.

## STAR Methods

### Mice

Male and female Kir6.2-deficient (Kir6.2–/–) and wildtype (WT; both B6C3F1/J mixed genetic background) were used in all experiments. Mice were group housed, given food and water ad libitum, and maintained on a 12:12 light/dark cycle. All procedures were carried out in accordance with an approved IACUC protocol from either Washington University in St. Louis, Wake Forest School of Medicine, or the University of Kentucky College of Medicine.

### 13C-glucose oral gavage and tissue collection for stable isotope resolved metabolomics

Kir6.2-/- and WT mice were single housed and fasted for 4 hours. Mice were given an oral gavage of 250 ul of 32mg/ml ^13^C-glucose (Cambridge Isotope Laboratories Inc., #110187-42-3) an estimated dose of 2mg/kg per animal for mice. 45 minutes post gavage, mice were cervically dislocated, and brain tissue was rapidly extracted. Samples were washed and flash frozen in liquid nitrogen. Stable isotope resolved metabolomics was performed as previously described (19).

### Electroencephalography/Electromyography Surgery

3-6 month old Kir6.2-/- and WT mice (N= 8-11 mice/genotype) were anesthetized with isoflurane and underwent stereotaxic surgery. Biosensor guide cannulas (MD-2255, BASi Research Products) were implanted bilaterally into the hippocampus (A/P -3mm, M/L +/- 3mm, D/V -1.8mm). Two stainless steel skull screws (MD-1310, BASi Research Products) were placed into the skull over the right frontal cortex (A/P 1mm, M/L -1mm) and left parietal cortex (A/P -2mm, M/L -1mm). These sites served as recording electrodes. A third reference electrode was placed over the cerebellum (A/P -6mm, M/L 0mm).

Insulated wire leads soldered to an EEG/EMG/Biosensor headmount (8402, Pinnacle Technology) and attached to the implanted skull screws for EEG recordings. A set of stainless steel wires were implanted into the nuchal muscles for EMG recordings. The screws, biosensor cannulas, EEG lead wires, and headmount were secured to the skull using acrylic dental cement (Reliance DuraLay). Post-surgery, mice were single housed and allowed to recover in recording cages (8288, Pinnacle Technology) placed into sound attenuating chambers (ENV-017M Med Associates). Mice were maintained on 12:12 light/dark cycle.

### Biosensor calibration

Biosensors were calibrated before insertion in mice per the manufacturer’s protocol (Pinnacle Technologies) and as described elsewhere (Naylor, 2012). Briefly, using the manufacturer’s calibration stage (7052, Pinnacle Technology), biosensors were inserted into 1xPBS at 37° C and allowed to stabilize for a baseline measurement. Two consecutive injections of the analyte of interest (L-Lactate or D-Glucose) were injected into this heated 1xPBS solution. A third interference analyte injections (ascorbic acid) was conducted. Finally, two additional consecutive injections of the analyte of interest were conducted. Biosensor readings were allowed to stabilize in between all injections. The resulting readout of all biosensors used in this experiment was strong, stepwise response to analytes of interest, and no response to the interferant analyte. The average change in response to each analyte injection was converted into a constant for that individual biosensor to be used for analyzing experimental data.

### EEG/EMG, glucose and lactate biosensor recording

72 hours post-surgery, mice were briefly anesthetized with isoflourane and two amperometric biosensors specific for either glucose or lactate (7004, Pinnacle Technology) were inserted into the guide cannula (n= 8-11 mice/genotype) into the left and right hippocampus. Biosensors were connected to a preamplifier (100x amplification, EEG high pass filter: 0.5 Hz, EMG high pass filter: 10 Hz (8406- 5SL, Pinnacle technology)) and inserted into the mouse’s headmount. The preamplifier was connected to the data acquisition system via a commutator (8401, Pinnacle Technology). Data sampling was transmitted at 250 Hz for EEG/EMG and 1 Hz for the biosensors. Mice were unanesthetized and freely allowed to move about their cages for the duration of recordings. Recordings began immediately.

Analysis was only performed after a stable baseline was reached for EEG/EMG and biosensors (Sirenia Acquisition software). 24-hour recordings of EEG/EMG and biosensor data taken from day 2 of the recording were used to establish a diurnal rhythm (n=8-11 mice per genotype).

### Biosensor Analysis

24 hours of biosensor data were analyzed from day 2 of recording. Nano-amp (nA) values per second were binned into 10 second averages. Using the calibration value for each biosensor, nA responses were converted into a concentration value of nano moles (mM) for each analyte of interest. The average concentration value over the 24h day was taken and used to create a percent of average concentration value for each 10 second bin. These 10 second percent of baseline values were then binned into 1 hour bins. Biosensors that exceeded a max change of 10% from baseline average across 24 hours were detrended using the detrend function in matlab to normalize data.

### Wake, NREM, and REM scoring

EEG/EMG data were exported and scored in 10-second epochs by a human trained in sleep scoring according to standard classifications (44) for wake (high frequency, low amplitude EEG, high amplitude EMG), NREM (low frequency, high amplitude EEG, low amplitude EMG), and REM (High frequency, low amplitude EEG, very low amplitude EMG) using Sirenia Sleep Software (n=8-11 mice per genotype) for 24 hours over the circadian day, beginning at lights on (ZT0).

### EEG power spectral analysis

EEG power spectral analysis was calculated using Sirenia Sleep Pro Software (Pinnacle Technology). A Fast Fourier transform (Hann window) was used to determine power spectra values for all epochs without artifacts and exported according to sleep and wake state. To determine absolute power, power spectra were binned in 1Hz bins and graphed for wake, NREM and REM. Relative power was generated by separating data into frequency bins in interest for sleep and wake behaviors for each 10s epoch (delta 0.5-4 Hz, theta 4-8 Hz, alpha 8-13 Hz, beta 13-30 Hz, gamma 30-50 Hz). A percentage of total power value was calculated for each frequency band and graphed according to sleep wake state.

### Sleep transition analysis

Sleep-wake transitions were evaluated for 2 hours post saline injection and, as, the EEG/EMG was dominated by wake and NREM bouts, REM epochs were excluded for this analysis (n=8-18 mice/genotype). Inclusion criteria for which sleep and wake transitions to include were as follows: NREM bouts lasting ≥ 5 minutes and Wake bouts lasting ≥ 10 minutes. These criteria were used to capture a transition that resulted in either sustained wake or NREM sleep, since mice do not have consolidated sleep. The 60 seconds preceding a transition were used as a baseline of ISF lactate levels. This baseline was used to calculate a percent change in ISF lactate in the 5 minutes following a NREM to wake or wake to NREM transition.

### Saline and glucose injections biosensor and sleep analysis protocol

Intraperitoneal (IP) injections were given to Kir6.2-/- mice only (n= 8) over a two-day period: saline (0.9% NaCl, 2g/kg) and glucose (50% dextrose, 2g/kg). Injections were given across the light period at specific times (Day 1: 10am and 3pm, Day 2: 10am) and the order of injections was randomized to avoid any circadian effect. Biosensor data was exported and converted into mM data in 10 seconds bins as described above. Baseline biosensor data was defined as data 10 minutes before each injection. Post-injection biosensor bins were compared to the 10-minute baseline and converted into a percentage change from baseline. % max change was determined by comparing % change from baseline of glucose injections relative to % change of baseline of saline injections. Changes in time spent in wake following the glucose injections were determined by comparing baseline % time in wake to % time in wake following each injection.

### Body composition and indirect calorimetry

3 days prior to indirect calorimetry, mice were single housed and underwent EchoMRI body composition analysis (EchoMRI). Mice were then transferred to the room containing Promethion Core metabolic system (Sable Systems International) and acclimated to room and single housing for 3 days while maintained on a 12:12 L:D cycle(40). Mice were then transferred into Promethion system for peripheral metabolic phenotyping (n=5/genotype) for 4 days. Day 2 of indirect calorimetry data was used for all analysis.

### Arousal challenge

3 month old WT and Kir6.2-/- mice (n = 3-4/group) were given an intraperitoneal injection of ketamine/xylazine (dose = 108.6mg/kg of ketamine, 16.8mg/kg of xylazine). Two independent investigators observed mice for a loss of righting relax. “Induction time” is the duration it took for the mice to lose their righting relax post-injection. “Emergence time” was recorded as the duration it took for a mouse to display the first signs of consciousness, including motor movement and a righting reflex, since losing their righting relax.

### Behavioral Testing

9-month-old female Kir6.2-/- and WT mice (N = 10-20 mice/genotype) were tested for behavioral differences in the Washington University Animal Behavior Core. Following 1 week habituation and handling, mice were evaluated for differences in locomotor activity and exploratory behavior, walking initiation, Morris Water Maze, and acoustic startle response. All tests were conducted during the light phase of the light/dark cycle, by a experimenter blind to the genotype of the mice.

### One-hour locomotor activity and exploratory behavior

To assess general activity levels and alterations in emotionality, mice were evaluated over a 1-h period in transparent (47.6 × 25.4 × 20.6 cm high) polystyrene enclosures. Each cage was surrounded by a frame containing a 4 × 8 matrix of photocell pairs, the output of which was fed to an on-line computer (Hamilton-Kinder, LLC, Poway, CA). The system software (Hamilton-Kinder, LLC) was used to define a 33 × 11 cm central zone and a peripheral or surrounding zone that was 5.5 cm wide with the sides of the cage being the outermost boundary. This peripheral area extended along the entire perimeter of the cage. Variables analyzed included the total number of ambulations and rearing on hindlimbs, as well as the number of entries, the time spent, and the distance traveled in the center area as well as the periphery surrounding the center.

### Walking Initiation

In the walking initiation test, mice were placed on a flat surface in the center of a 21 cm x 21 cm square. The amount of time the mouse took to leave the square was recorded as a measure of initiation of movement.

### Morris water maze

Water maze testing was conducted as previously described (Yuede et al., 2021). Briefly, cued, place and probe trials were conducted in a galvanized steel pool, measuring 120 cm in diameter, and filled with opaque water (diluted nontoxic white tempera paint). The PVC escape platform measured 11.5 cm in diameter. A digital video camera connected to a PC computer with the computer software program ANY-maze (Stoelting Co.) tracked swimming pathway of the mouse to the escape platform and quantified path length, latency to find escape platform, and swimming speeds. On two consecutive days, animals received four cued trials to habituate to the swimming task procedure and control for any differences in swimming, visual, or motivational performance in the test. A red tennis ball atop a rod was attached to the escape platform and served as a visual cue for the platform. To prevent spatial learning, the escape platform was moved to a different quadrant location for each trial. The mouse was released from the quadrant opposite to the platform location and allowed 60s to locate the platform. Once the mouse found the platform, it was allowed to remain there for 10s before being returned to its home cage. Three days following visible platform testing, the cue was removed from the platform, and it was submerged 1 cm under the water in a fixed location for the hidden platform tests to evaluate spatial learning. Animals received two blocks of two consecutive trials on five consecutive days, with an inter-trial interval between 30–90s and approximately 2 hr separating trial blocks. The escape platform remained in the same quadrant location for all trials and distal cues were placed on the walls of the room to support spatial learning. The mouse was released from a different location for each trial on each day. The mouse was allowed 60s to find the escape platform and allowed to sit on it for 10s before being returned to its home cage. Cued and hidden platform trials were combined into blocks of two or four trials for analyses, respectively. One hour following completion of hidden platform trials on the 5th day of training, the escape platform was removed from the pool and one 60s probe trial was conducted to assess memory retention for the location of the platform.

### Acoustic Startle Response/Prepulse Inhibition

Startle response to a 120 dB auditory stimulus pulse (40 ms broadband burst) and PPI (response to a prepulse plus the startle pulse) were measured concurrently in mice using the Kinder Scientific Startle Reflex system (Poway, CA). Beginning at stimulus onset, 1 ms force readings were averaged to obtain an animal’s startle amplitude. A total of 20 startle trials were presented over a 20 min test period during which the first 5 min served as an acclimation period when no stimuli above the 70 dB white noise background were presented. The session began and ended by presenting 5 consecutive startle (120 db pulse alone) trials unaccompanied by other trial types. The middle 10 startle trials were interspersed with PPI trials (consisting of an additional 30 presentations of 120 dB startle stimuli preceded by pre- pulse stimuli of either 4, 12, or 20 dB above background (10 trials for each PPI trial type). A percent PPI score for each trial was calculated using the following equation: %PPI= 100*(ASRstartle pulse alone - ASRprepulse+startle pulse)/ASRstartle pulse alone.

### Jonckheere–Terpstra–Kendall (JTK) analysis

Data was preprocessed to average group values for each ZT timepoint. JTK analysis was done using R package MetaCycle. Adjusted p-value was reported to determine the significance of rhythmicity. A p-value of p<0.05 was used as a cutoff for determining statistical significance.

## Quantification and statistical analysis

All EEG and sleep data was scored using the Sirenia Sleep Pro (Pinnacle Technologies) and exported for analysis. All of the statistical analysis was done using GraphPad Prism 10 (GraphPad Software, LLC). All data is reported as means with +/- SEM. All details of statistical tests and corrections, subject numbers, and p-values are located in the figure legends. A p-value of p<0.05 was used as a cutoff for determining statistical significance. Specific statistical tests used were as follows:

Figure 1: 2-way ANOVA with Tukey multiple comparisons correction; unpaired t-test

Figure 2: 2-way ANOVA with Sidak or Bonferroni multiple comparisons correction; unpaired t-test

Figure 3: 2-way ANOVA with Tukey multiple comparisons correction

Figure 4: 2-way ANOVA with Sidak or Tukey multiple comparisons correction for post hoc analysis; One-way ANOVA with Dunnett multiple comparisons correction for post hoc analysis

Table 1: Jonckheere–Terpstra–Kendall analysis Figures S1 & S2: Unpaired t-test

Figures S3-S5: Pearson’s R correlation

## Author Contributions

SLM, NJC, and CC conceived of the study. SLM, NJC, CC, ESM, and LAJ contributed to study design. NCJ, CC, HCW, CMY, PWS, and JAS performed experiments. SLM, NJC, CC, HCW, CMY, PWS, and LAJ performed data analysis and data interpretation. NJC and SLM wrote the manuscript. All authors discussed the results and commented on the manuscript.

## Acknowledgements

We would like to thank Jamie Hicks, MA for involvement in behavior data collection. We would like to thank Drs. Robert Gould and Marc Raichle for intellectual discussion related to this manuscript. We would like to acknowledge the following grants: K01AG050719 (SLM), R01AG068330 (SLM), BrightFocus Foundation A20201775S (SLM), T32NS115704 (NJC), F31AG066302 (CC),

R01AG060056 (LAJ), R01AG062550 (LAJ), R01AG080589 (LAJ), and the Alzheimer’s Association (LAJ). Research reported in this publication was supported by an Institutional Development Award (IDeA) from the National Institute of General Medical Sciences of the National Institutes of Health under grant number P30GM127211 and the NIH Center of Biomedical Research Excellence (COBRE) in CNS Metabolism (CNS-Met) under grant number P20GM148326. The Washington University Animal Behavior Core is supported in part by funds from the McDonnell Center for Systems Neuroscience, the McDonnell Center for Cellular and Molecular Neuroscience, and the Taylor Family Institute.

---

**Supplemental figure 1.**
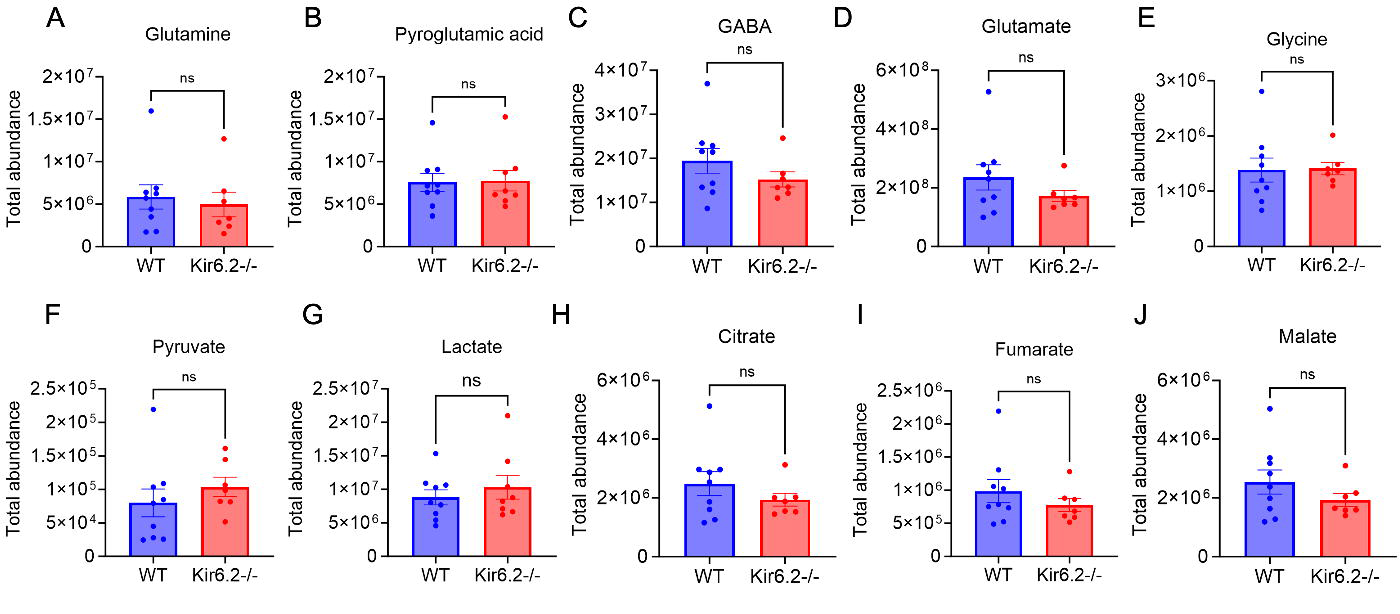
Brain metabolite total abundance quantified by stable isotope resolved metabolomics. (A-J) Total abundance is unaltered across groups. Data reported as means± SEM. n = 8-9 mice/genotype. Significance determined via unpaired t-test.

**Supplemental figure 2.**
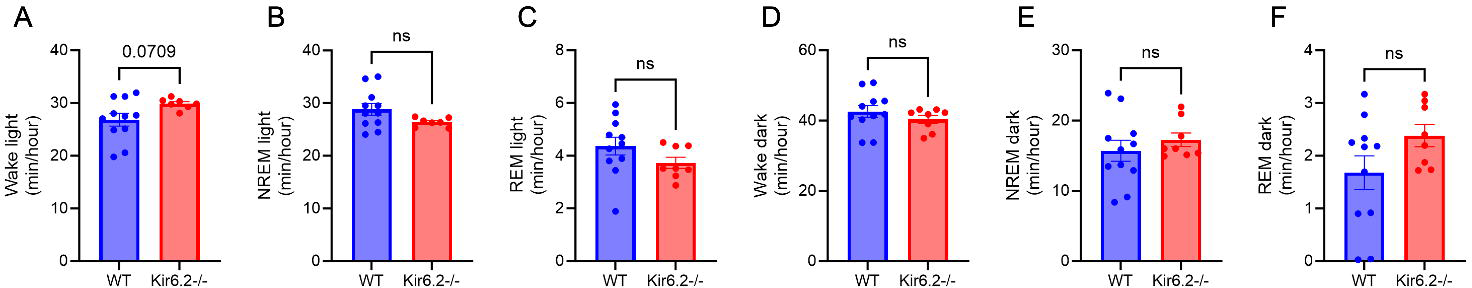
Time spent in sleep-wake states across light and dark periods. (A) Kir6.2-/- mice trend towards increased time spent in wake during the light period (p=0.0709). (B-C) There is no change in time spent in NREM and REM sleep in Kir6.2-/- mice during the light period. (D-F) No change in time spent in wake, NREM, or REM sleep in Kir6.2-/- mice during the dark period. Data reported as means± SEM. n = 9-10 mice/genotype. Minute/hour sleep-wake state significance determined by unpaired t-tests.

**Supplemental figure 3.**
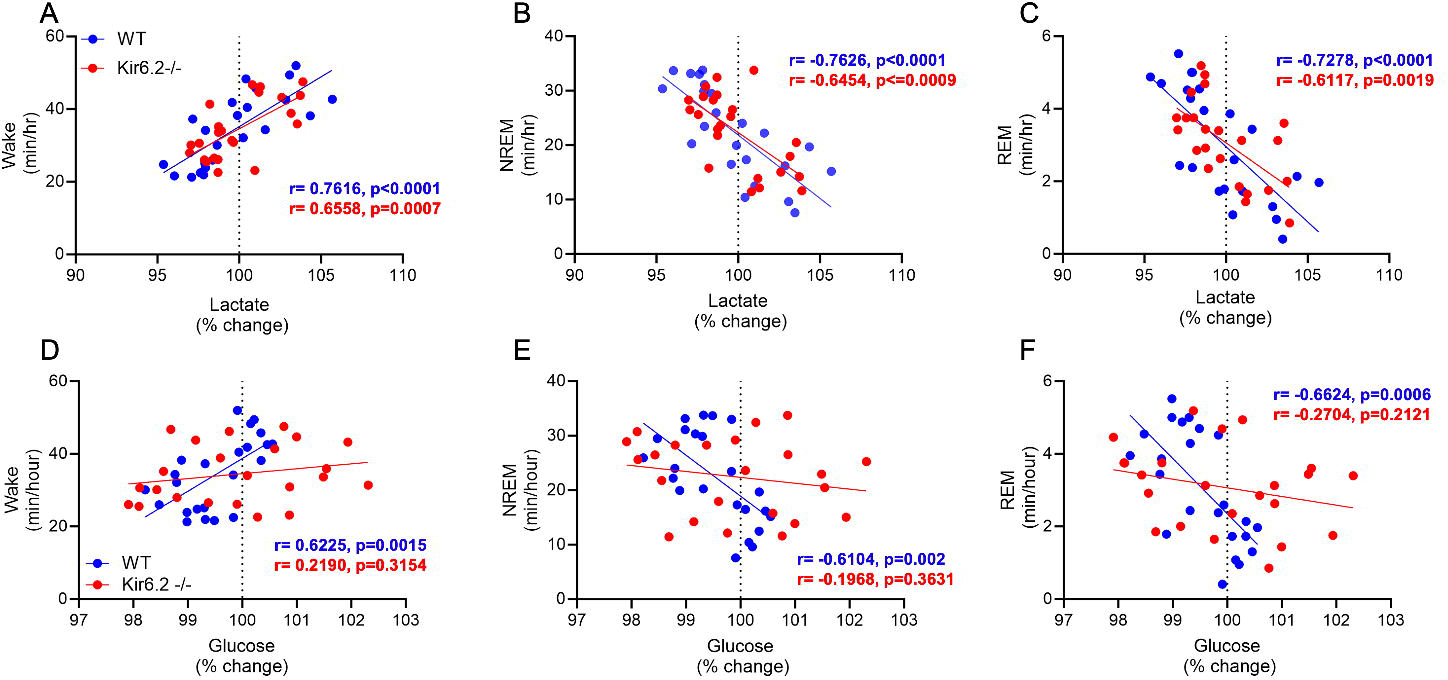
Correlations of brain ISF lactate and ISF glucose with sleep and wake states across the diurnal day. (A) ISF lactate is positively correlated with increases in time spent in wake in Kir6.2-/- and WT mice. (B-C) In Kir6.2--/- and WT mice, ISF lactate is negatively correlated with NREM and REM sleep. (D) ISF glucose is positively correlated with wake in WT but not Kir6.2-/- mice. (E-F) ISF glucose is negatively correlated with time spent in NREM and REM sleep in WT but not Kir6.2-/- mice. Data reported as means± SEM. n = 8- 11 mice/genotype. Significance determined by Pearson’s R correlation.

**Supplemental figure 4.**
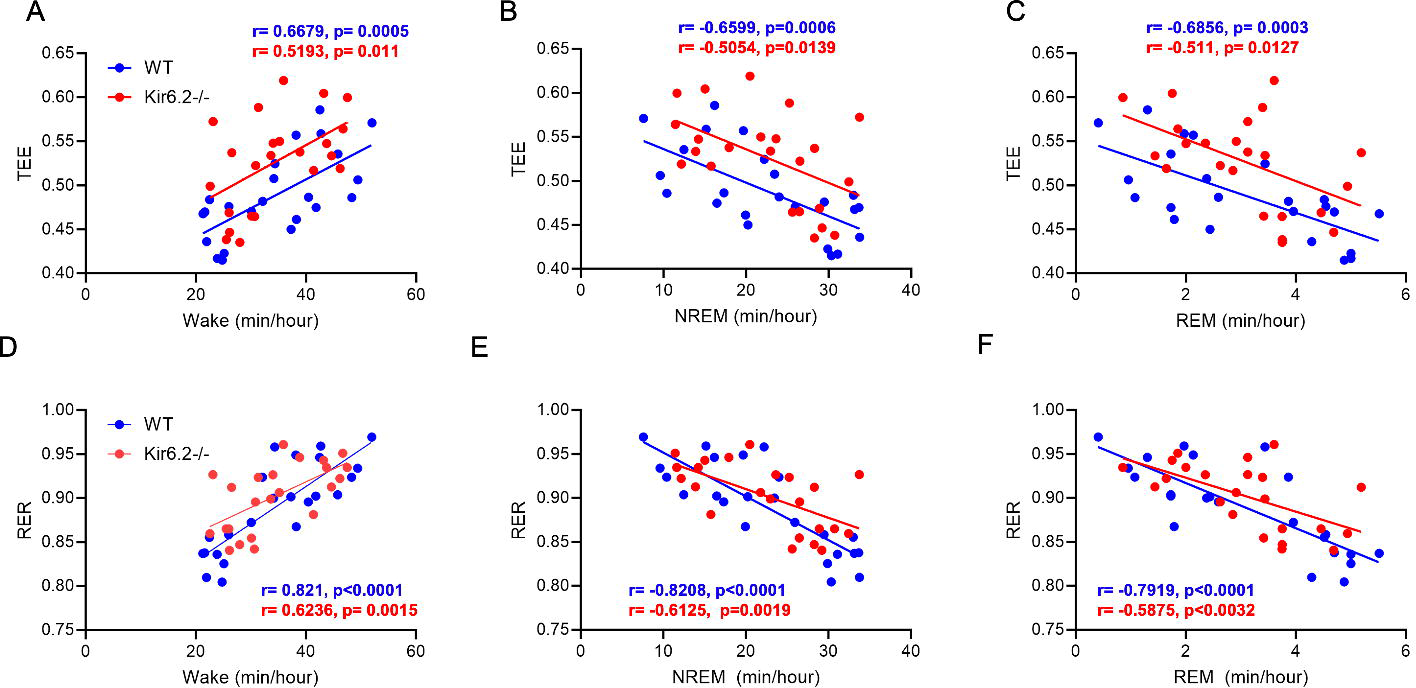
TEE and RER are correlated with sleep and wake states. (A-C) TEE is positively correlated with time spent in wake in both Kir6.2-/- and WT mice, and negatively correlated with NREM and REM sleep. (D-E) RER is positively correlated with time spent in wake in both Kir6.2-/- and WT mice, and negatively correlated with NREM and REM sleep. Data reported as means± SEM. n = 5-11 mice/genotype. Significance determined by Pearson’s R correlation.

**Supplemental figure 5.**
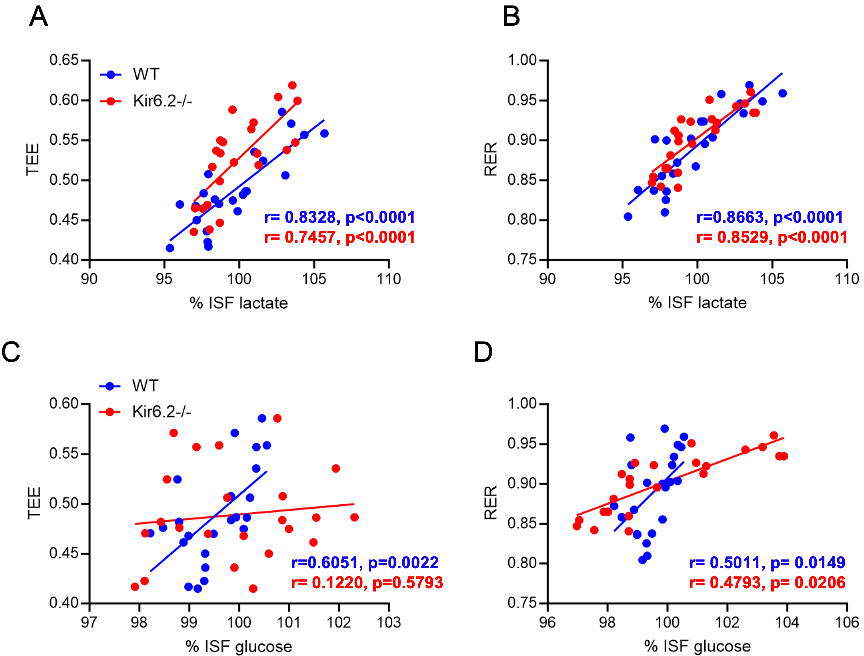
The relationship of peripheral and central metabolism. (A-B) TEE and RER are positively correlated with brain ISF lactate in both Kir6.2-/- and WT mice. (C) Kir6.2 -/- mice lose relationship between TEE and brain ISF glucose seen in WT mice. (D) Both Kir6.2-/- and WT mice RER are positively correlated with brain ISF glucose. Data reported as means± SEM. n = 5-11 mice/genotype. Significance determined by Pearson’s R correlation.

**Supplemental Table 1.**
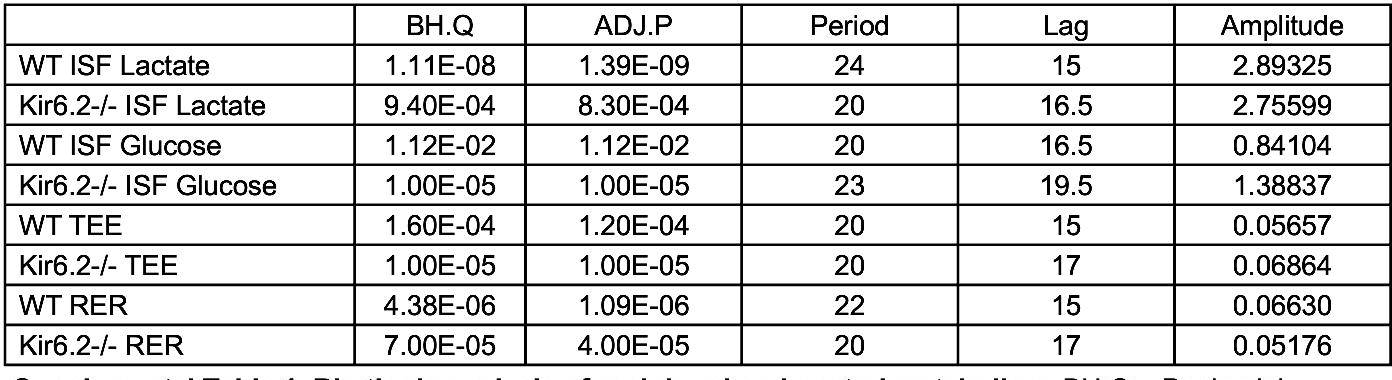
Rhythmic analysis of peripheral and central metabolism. BH.Q = Benjamini­ Hochberg procedure. ADJ.P = adjusted P value. Period = length of cycle. LAG = time of peak expression. Amplitude = peak expression. JTK analysis was performed for each genotype and condition.

|  | BH.Q | ADJ.P | Period | Lag | Amplitude |
| --- | --- | --- | --- | --- | --- |
| WT ISF Lactate | 1.11E-08 | 1.39E-09 | 24 | 15 | 2.89325 |
| Kir6.2-/- ISF Lactate | 9.40E-04 | 8.30E-04 | 20 | 16.5 | 2.75599 |
| WT ISF Glucose | 1.12E-02 | 1.12E-02 | 20 | 16.5 | 0.84104 |
| Kir6.2-/- ISF Glucose | 1.00E-05 | 1.00E-05 | 23 | 19.5 | 1.38837 |
| WT TEE | 1.60E-04 | 1.20E-04 | 20 | 15 | 0.05657 |
| Kir6.2-/- TEE | 1.00E-05 | 1.00E-05 | 20 | 17 | 0.06864 |
| WT RER | 4.38E-06 | 1.09E-06 | 22 | 15 | 0.06630 |
| Kir6.2-/- RER | 7.00E-05 | 4.00E-05 | 20 | 17 | 0.05176 |

## References

1. Aalling NN, Nedergaard M, and DiNuzzo M. Cerebral Metabolic Changes During Sleep. Curr Neurol Neurosci Rep. 2018;18(9):57.

2. DiNuzzo M, and Nedergaard M. Brain energetics during the sleep-wake cycle. Curr Opin Neurobiol. 2017;47:65–72.

3. Rempe MJ, and Wisor JP. Cerebral lactate dynamics across sleep/wake cycles. Front Comput Neurosci. 2014;8:174.

4. Naylor E, Aillon DV, Barrett BS, Wilson GS, Johnson DA, Johnson DA, et al. Lactate as a biomarker for sleep. Sleep. 2012;35(9):1209–22.

5. Carroll CM, and Macauley SL. The Interaction Between Sleep and Metabolism in Alzheimer’s Disease: Cause or Consequence of Disease? Front Aging Neurosci. 2019;11:258.

6. Carroll CM, Stanley M, Raut RV, Constantino NJ, Irmen RE, Mitra A, et al. Acute hyper- and hypoglycemia uncouples the metabolic cooperation between glucose and lactate to disrupt sleep. bioRxiv. 2022.

7. Zuend M, Saab AS, Wyss MT, Ferrari KD, Hosli L, Looser ZJ, et al. Arousal-induced cortical activity triggers lactate release from astrocytes. Nat Metab. 2020;2(2):179–91.

8. Marcheva B, Ramsey KM, Peek CB, Affinati A, Maury E, and Bass J. Circadian clocks and metabolism. Handb Exp Pharmacol. 2013(217):127–55.

9. Karagiannis A, Gallopin T, Lacroix A, Plaisier F, Piquet J, Geoffroy H, et al. Lactate is an energy substrate for rodent cortical neurons and enhances their firing activity. Elife. 2021;10.

10. Grizzanti J, Moritz WR, Pait MC, Stanley M, Kaye SD, Carroll CM, et al. KATP channels are necessary for glucose-dependent increases in amyloid-beta and Alzheimer’s disease-related pathology. JCI Insight. 2023;8(10).

11. Burdakov D, Luckman SM, and Verkhratsky A. Glucose-sensing neurons of the hypothalamus. Philos Trans R Soc Lond B Biol Sci. 2005;360(1464):2227-35.

12. Parsons MP, and Hirasawa M. ATP-sensitive potassium channel-mediated lactate effect on orexin neurons: implications for brain energetics during arousal. J Neurosci. 2010;30(24):8061–70.

13. Clasadonte J, Scemes E, Wang Z, Boison D, and Haydon PG. Connexin 43-Mediated Astroglial Metabolic Networks Contribute to the Regulation of the Sleep-Wake Cycle. Neuron. 2017;95(6):1365–80 e5.

14. Nelson PT, Jicha GA, Wang WX, Ighodaro E, Artiushin S, Nichols CG, and Fardo DW. ABCC9/SUR2 in the brain: Implications for hippocampal sclerosis of aging and a potential therapeutic target. Ageing Res Rev. 2015;24(Pt B):111–25.

15. Allebrandt KV, Amin N, Muller-Myhsok B, Esko T, Teder-Laving M, Azevedo RV, et al. A K(ATP) channel gene effect on sleep duration: from genome-wide association studies to function in Drosophila. Mol Psychiatry. 2013;18(1):122–32.

16. Scheinfeldt LB, Gharani N, Kasper RS, Schmidlen TJ, Gordon ES, Jarvis JP, et al. Using the Coriell Personalized Medicine Collaborative Data to conduct a genome-wide association study of sleep duration. Am J Med Genet B Neuropsychiatr Genet. 2015;168(8):697–705.

17. Ota H, Tamaki S, Itaya-Hironaka A, Yamauchi A, Sakuramoto-Tsuchida S, Morioka T, et al. Attenuation of glucose-induced insulin secretion by intermittent hypoxia via down-regulation of CD38. Life Sci. 2012;90(5-6):206–11.

18. Landmeier KA, Lanning M, Carmody D, Greeley SAW, and Msall ME. ADHD, learning difficulties and sleep disturbances associated with KCNJ11-related neonatal diabetes. Pediatr Diabetes. 2017;18(7):518–23.

19. Williams HC, Piron MA, Nation GK, Walsh AE, Young LEA, Sun RC, and Johnson LA. Oral Gavage Delivery of Stable Isotope Tracer for In Vivo Metabolomics. Metabolites. 2020;10(12).

20. Pitchford B, and Arnell KM. Resting EEG in alpha and beta bands predicts individual differences in attentional breadth. Conscious Cogn. 2019;75:102803.

21. Foster JJ, Sutterer DW, Serences JT, Vogel EK, and Awh E. Alpha-Band Oscillations Enable Spatially and Temporally Resolved Tracking of Covert Spatial Attention. Psychol Sci. 2017;28(7):929–41.

22. Brancaccio A, Tabarelli D, Bigica M, and Baldauf D. Cortical source localization of sleep-stage specific oscillatory activity. Sci Rep. 2020;10(1):6976.

23. Cantero JL, Atienza M, and Salas RM. Human alpha oscillations in wakefulness, drowsiness period, and REM sleep: different electroencephalographic phenomena within the alpha band. Neurophysiol Clin. 2002;32(1):54–71.

24. Mizuseki K, and Buzsaki G. Theta oscillations decrease spike synchrony in the hippocampus and entorhinal cortex. Philos Trans R Soc Lond B Biol Sci. 2014;369(1635):20120530.

25. Buzsaki G. Theta oscillations in the hippocampus. Neuron. 2002;33(3):325–40.

26. Snipes S, Krugliakova E, Meier E, and Huber R. The Theta Paradox: 4-8 Hz EEG Oscillations Reflect Both Sleep Pressure and Cognitive Control. J Neurosci. 2022;42(45):8569–86.

27. Vyazovskiy VV, and Tobler I. Theta activity in the waking EEG is a marker of sleep propensity in the rat. Brain Res. 2005;1050(1-2):64–71.

28. Dadashi M, Birashk B, Taremian F, Asgarnejad AA, and Momtazi S. Effects of Increase in Amplitude of Occipital Alpha & Theta Brain Waves on Global Functioning Level of Patients with GAD. Basic Clin Neurosci. 2015;6(1):14–20.

29. Castegnetti G, Bush D, and Bach DR. Model of theta frequency perturbations and contextual fear memory. Hippocampus. 2021;31(4):448–57.

30. Goodman MS, Kumar S, Zomorrodi R, Ghazala Z, Cheam ASM, Barr MS, et al. Theta-Gamma Coupling and Working Memory in Alzheimer’s Dementia and Mild Cognitive Impairment. Front Aging Neurosci. 2018;10:101.

31. Moretti DV, Prestia A, Binetti G, Zanetti O, and Frisoni GB. Increase of theta frequency is associated with reduction in regional cerebral blood flow only in subjects with mild cognitive impairment with higher upper alpha/low alpha EEG frequency power ratio. Front Behav Neurosci. 2013;7:188.

32. Thomas R, Zimmerman SD, Yuede KM, Cirrito JR, Tai LM, Timson BF, and Yuede CM. Exercise Training Results in Lower Amyloid Plaque Load and Greater Cognitive Function in an Intensity Dependent Manner in the Tg2576 Mouse Model of Alzheimer’s Disease. Brain Sci. 2020;10(2).

33. McKenna JT, Tartar JL, Ward CP, Thakkar MM, Cordeira JW, McCarley RW, and Strecker RE. Sleep fragmentation elevates behavioral, electrographic and neurochemical measures of sleepiness. Neuroscience. 2007;146(4):1462–73.

34. Huber R, Ghilardi MF, Massimini M, Ferrarelli F, Riedner BA, Peterson MJ, and Tononi G. Arm immobilization causes cortical plastic changes and locally decreases sleep slow wave activity. Nat Neurosci. 2006;9(9):1169–76.

35. Werth E, Dijk DJ, Achermann P, and Borbely AA. Dynamics of the sleep EEG after an early evening nap: experimental data and simulations. Am J Physiol. 1996;271(3 Pt 2):R501-10.

36. Borbely AA, and Achermann P. Concepts and models of sleep regulation: an overview. J Sleep Res. 1992;1(2):63–79.

37. Valderrama M, Crepon B, Botella-Soler V, Martinerie J, Hasboun D, Alvarado-Rojas C, et al. Human gamma oscillations during slow wave sleep. PLoS One. 2012;7(4):e33477.

38. Hutchison IC, and Rathore S. The role of REM sleep theta activity in emotional memory. Front Psychol. 2015;6:1439.

39. Wisor JP, Rempe MJ, Schmidt MA, Moore ME, and Clegern WC. Sleep slow-wave activity regulates cerebral glycolytic metabolism. Cereb Cortex. 2013;23(8):1978–87.

40. Farmer BC, Williams HC, Devanney NA, Piron MA, Nation GK, Carter DJ, et al. APOEpsilon4 lowers energy expenditure in females and impairs glucose oxidation by increasing flux through aerobic glycolysis. Mol Neurodegener. 2021;16(1):62.

41. Foley J, Blutstein T, Lee S, Erneux C, Halassa MM, and Haydon P. Astrocytic IP(3)/Ca(2+) Signaling Modulates Theta Rhythm and REM Sleep. Front Neural Circuits. 2017;11:3.

42. Platt B, and Riedel G. The cholinergic system, EEG and sleep. Behav Brain Res. 2011;221(2):499-504.

43. Oken BS, Salinsky MC, and Elsas SM. Vigilance, alertness, or sustained attention: physiological basis and measurement. Clin Neurophysiol. 2006;117(9):1885–901.

44. Moser D, Anderer P, Gruber G, Parapatics S, Loretz E, Boeck M, et al. Sleep classification according to AASM and Rechtschaffen & Kales: effects on sleep scoring parameters. Sleep. 2009;32(2):139–49.

